# Network-based prediction and functional validation of metformin for potential treatment of atrial fibrillation using human inducible pluripotent stem cell-derived atrial-like cardiomyocytes

**DOI:** 10.1101/2021.09.17.460826

**Authors:** Jessica Castrillon Lal, Yadi Zhou, Shamone R. Gore-Panter, Julie H. Rennison, John Barnard, David R. Van Wagoner, Feixiong Cheng, Mina K. Chung

## Abstract

Atrial fibrillation (AF) is a significant cause of morbidity and mortality, and effective therapeutic interventions are lacking. Here, we harness an integrative, network medicine approach to repurpose FDA-approved drugs for AF. We hypothesize that the use of an unbiased method for prioritizing AF drugs using patient transcriptomics data can help to identify alternative therapeutic strategies and mechanism-of-action for these drugs. To achieve this, we first characterized the molecular networks specific to AF by incorporating transcriptomic data of left atrial tissue. We quantified the network proximity of genes differentially expressed in AF to drug targets to identify putative drugs for repurposing. We identified nine high-confidence drug candidates that were validated using enrichment analysis of drug-gene signatures in human cell lines. We identified metformin for the potential treatment of AF and validated its use in human inducible pluripotent stem cell-derived atrial-like cardiomyocytes. We identified AF-specific dysregulated networks enriched in cardiac metabolism, ion transport, and immune pathways that were improved following metformin treatment. In summary, this study utilized network-based approaches for rapid identification of drugs that may be repurposed for AF treatment and validated metformin as a candidate drug using a robust human atrial cell model.

## Introduction

Atrial fibrillation (AF) is the most prevalent cardiac arrhythmia in the western world, affecting 1% to 2% of the general population ^1–3^. The incidence in the US, currently estimated at 5-6 million individuals, is expected to increase to 12 million by 2030 ^4^. As AF is silent in 5-35% of diagnosed patients, the prevalence is likely higher. While the early stages of AF are considered benign, ^5^ persistent and long-standing persistent forms of AF are associated with a substantial increase in mortality with a 1.5 and 1.9 fold increase in the hazard ratio in men and women, respectively ^6^. AF is also associated with a higher risk of stroke, heart failure, and dementia ^7, 8^. AF patients are 30% more likely to be hospitalized at least once annually, resulting in a significant financial burden to patients and the healthcare system ^9^.

Clinical management of AF requires improvement. The primary goals of AF clinical management are controlling heart rate, restoring and maintaining sinus rhythm, and preventing thromboembolic. Drugs for controlling the ventricular rate and restoring sinus rhythm include β-adrenergic blockers, cardiac glycosides, and ion channel blockers; these drugs have potential side effects that include life-threatening proarrhythmic. Clinical management of AF is further limited by high toxicity rates of anti-arrhythmic drugs ^10–12^. AF ablation, including pulmonary vein isolation and/or atrial substrate ablation, are invasive strategies with limited success and can also be associated with major complications ^13^, ^14^. Anticoagulants such as direct oral anticoagulants or warfarin and/or left atrial appendage closure are used to prevent thromboembolism ^8^. Over 10 years have passed since the last antiarrhythmic drugs were approved for AF. Novel network medicine approaches may enable the identification of drug targets for AF that may be less toxic to patients using drug repurposing strategies ^15, 16^. Recent advances in network medicine and systems biology have enabled novel approaches for drug discovery or drug repurposing ^16–19^. Pathways for AF treatment can include multiple targets that share biological networks. This approach allows for discovering novel drug targets and minimizes the risk of toxicity by considering all drug-protein interactions within a protein-protein network ^15, 20^. Integrative analyses of genomics, transcriptomics, and interactomics (protein-protein interactions [PPIs]) have not yet been fully exploited for AF drug repurposing. Here, we used network-based approaches to prioritize drug repurposing for AF by unique integration of transcriptomics data from left atrial tissues and drug-induced gene signatures from human inducible pluripotent stem cell-derived atrial-like cardiomyocytes under the human protein-protein interactome (**Figure 1**).

**Figure 1.**
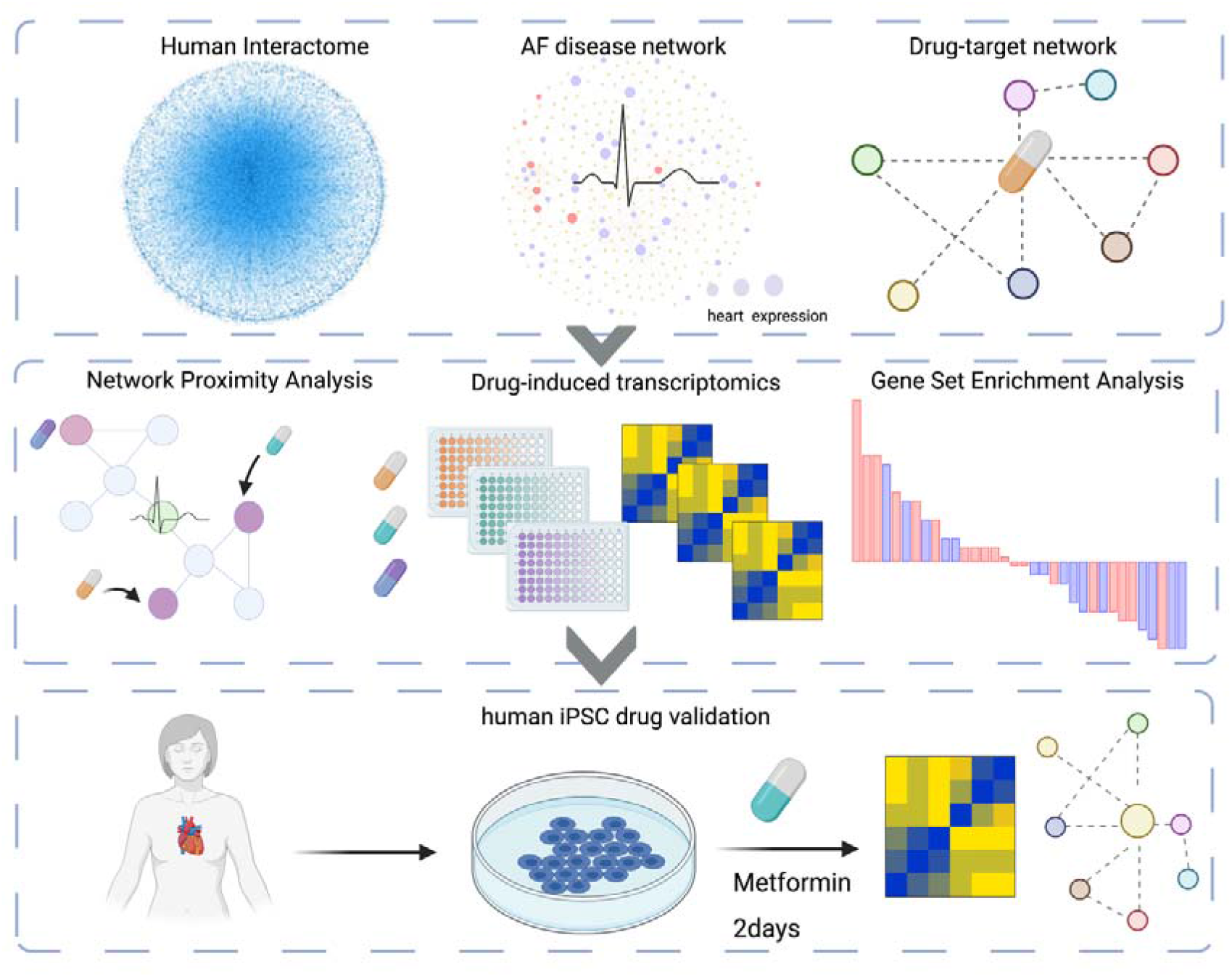
Study overview. Here we utilize systems pharmacology-based network medicine platform to quantify the proximity of interaction between the atrial fibrillation (AF) interactome nodes and drug targets in the human PPI network. Gene set enrichment analysis of known targets are used to validate our *in silico* approach and nominate candidate drugs. Using human derived induced pluripotent stem cells from AF patients, we were able to validate metformin as a repurposed drug for AF.

## Results

We characterized molecular networks specific to AF by incorporating transcriptomic data of genes differentially expressed in AF versus normal sinus rhythm of the left atrial appendage (LAA) tissues collected from 251 patients who underwent elective cardiac surgery to treat AF, valve disease, or other cardiac disorder (**Table 1**) ^21^. We further applied network-based discovery to identify drug candidates specific to AF. We demonstrate that drugs that can reverse AF-dysregulated gene expression may preserve tissue viability (**Figure 1B**) and help to identify alternative treatment strategies (**Figure 1C**).

**Table 1.**
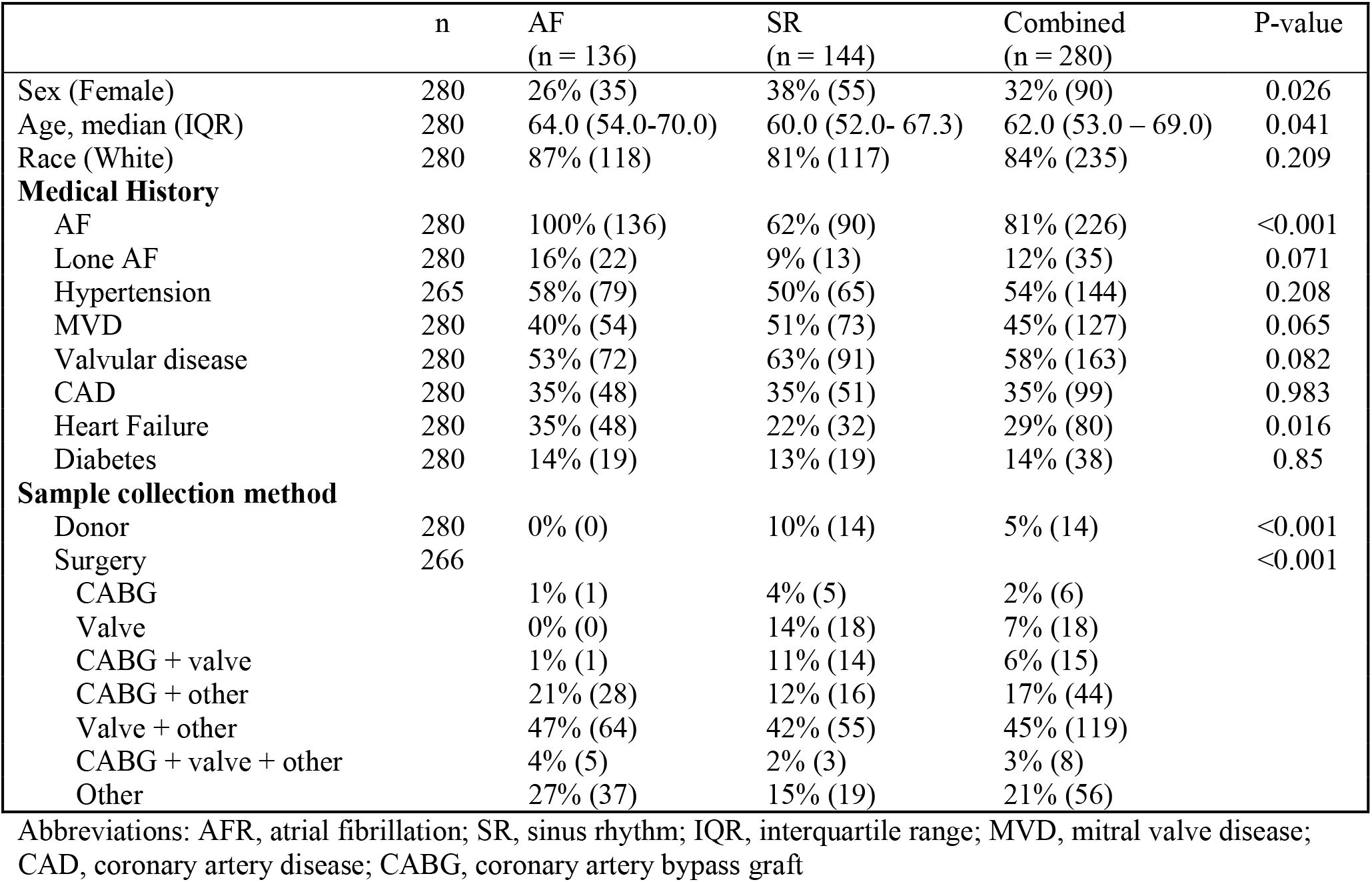
Human subject demographics.

|  | n | AF<br>(n = 136) | SR<br>(n = 144) | Combined<br>(n = 280) | P-value |
| --- | --- | --- | --- | --- | --- |
| Sex (Female) | 280 | 26% (35) | 38% (55) | 32% (90) | 0.026 |
| Age, median (IQR) | 280 | 64.0 (54.0-70.0) | 60.0 (52.0- 67.3) | 62.0 (53.0 – 69.0) | 0.041 |
| Race (White) | 280 | 87% (118) | 81% (117) | 84% (235) | 0.209 |
| **Medical History** |  |  |  |  |  |
| AF | 280 | 100% (136) | 62% (90) | 81% (226) | <0.001 |
| Lone AF | 280 | 16% (22) | 9% (13) | 12% (35) | 0.071 |
| Hypertension | 265 | 58% (79) | 50% (65) | 54% (144) | 0.208 |
| MVD | 280 | 40% (54) | 51% (73) | 45% (127) | 0.065 |
| Valvular disease | 280 | 53% (72) | 63% (91) | 58% (163) | 0.082 |
| CAD | 280 | 35% (48) | 35% (51) | 35% (99) | 0.983 |
| Heart Failure | 280 | 35% (48) | 22% (32) | 29% (80) | 0.016 |
| Diabetes | 280 | 14% (19) | 13% (19) | 14% (38) | 0.85 |
| **Sample collection method** |  |  |  |  |  |
| Donor | 280 | 0% (0) | 10% (14) | 5% (14) | <0.001 |
| Surgery | 266 |  |  |  | <0.001 |
| CABG |  | 1% (1) | 4% (5) | 2% (6) |  |
| Valve |  | 0% (0) | 14% (18) | 7% (18) |  |
| CABG + valve |  | 1% (1) | 11% (14) | 6% (15) |  |
| CABG + other |  | 21% (28) | 12% (16) | 17% (44) |  |
| Valve + other |  | 47% (64) | 42% (55) | 45% (119) |  |
| CABG + valve + other |  | 4% (5) | 2% (3) | 3% (8) |  |
| Other |  | 27% (37) | 15% (19) | 21% (56) |  |

Abbreviations: AFR, atrial fibrillation; SR, sinus rhythm; IQR, interquartile range; MVD, mitral valve disease; CAD, coronary artery disease; CABG, coronary artery bypass graft

### Atrial fibrillation disease network

We generated an integrative network-based approach to identify molecular networks of AF. We hypothesize that this network-medicine approach will enhance candidate drug prioritization for the treatment of AF. In prior work, we describe a structurally resolved human protein-protein interactome generated from five types of PPI ^16, 18, 22-24^. To depict the human interactome network, we gathered associated proteins from various bioinformatic and systems biology databases across 5 levels of evidence: (i) binary PPIs tested by high-throughput yeast two-hybrid systems (Y2H); (ii) binary, physical PPIs from protein 3D structures; (iii) kinase-substrate interactions identified in the literature; (iv) signaling networks identified in the literature; and (v) literature-curated PPIs identified by affinity purification followed by mass spectrometry, Y2H, or found in the literature. To depict the AF interactome, we gathered RNA-sequencing (RNASeq) data from a biobank of left atrial tissues obtained from cardiac surgery patients at the Cleveland Clinic to identify testable AF disease modules under the human proteinprotein interactome network model. Here, we identified 491 differentially expressed genes in AF cases (i.e., hearts in atrial fibrillation at the time of acquisition) compared to sinus rhythm cases (adjusted P-value [q] < 0.05). The AF disease module (defined by the largest connected component in the human interactome) shown in **Figure 2** includes 245 unique proteins (nodes) and 350 PPIs (edges). We identified several AF-specific hub proteins related to cardiac integrity and metabolic fitness, including *LDHB, CDH2, UQCRH, PDLIM5, COPS5*, and *OXCT1*. Gene Ontology (GO) enrichment analysis identified the following biological processes significant in AF differentially expressed genes: the endoplasmic reticulum unfolded protein response, calcium and potassium ion transport, succinyl-CoA metabolism, collagen biosynthesis, membrane repolarization, and cardiac muscle relaxation (**Suppl. Figure 1A**). Similarly, KEGG pathway enrichment analysis identified propanoate metabolism, proteasome, regulated calcium reabsorption, and cardiac muscle contraction as pathways significantly enriched in AF (**Suppl. Figure 1B**).

**Figure 2.**
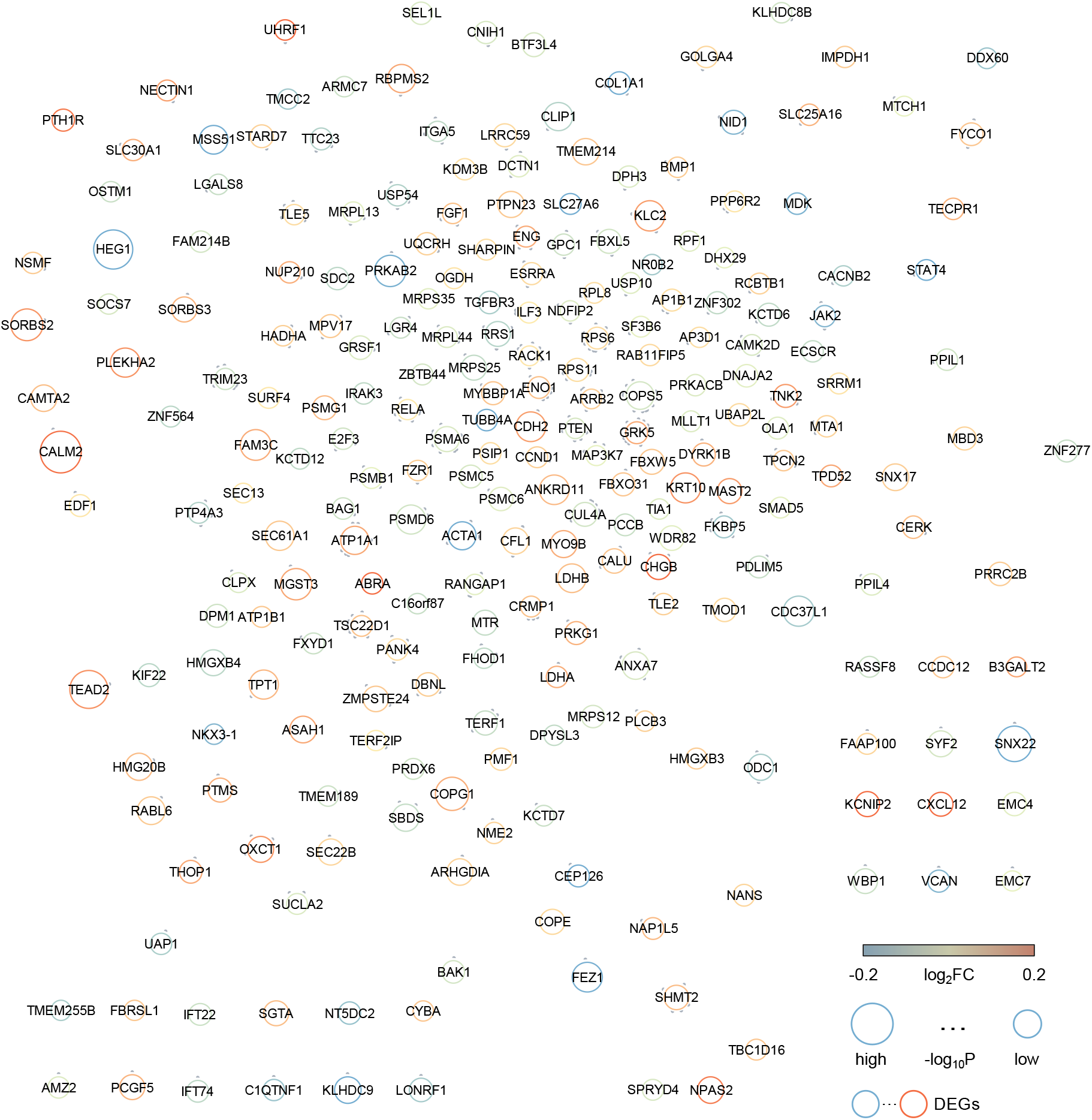
The atrial fibrillation interactome. The network highlights the atrial fibrillation (AF) interactome connects 245 AF enriched genes. Node size is proportional to -log_10_P-value, and color corresponds to log_2_ fold change (log_2_FC) in AF compared to normal sinus rhythm (see Methods).

### Network-based drug repurposing for AF

We harnessed drug-target interactions from 3,000 FDA-approved or clinically investigational drugs obtained from the Drugbank database to assemble the drug-target network (**Figure 1B**) ^25^. The network proximity of drug targets to AF disease module was calculated using Jaccard Index and Overlap coefficient (see Methods). To correct literature bias of well-studied proteins in the human interactome, we normalized calculated values to Z-score values and performed 1,000 permutation tests as described in prior work ^23, 24^. Using a Z-score cutoff value Z < −1, we focused on 55 drug candidates from total 2,891 drugs (**Figure 3A**). To narrow our repurposed drug list further, we used gene expression data of drug-treated human cell lines from the Connectivity Map database to prioritize drug candidates ^26^. Gene set enrichment analysis (GSEA) was performed and enrichment score (ES) was calculated to nominate drug candidates using a cutoff of ES > 0 and P <0.05. Here we identified 9 potential candidates: phenformin [Z= −2.475], metformin [Z= −1.97], furosemide [Z= −1.912], indapamide [Z= −1.691], metacycline [Z= −1.674], rofecoxib [Z= −1.67], dantrolene [Z= - 1.193], alclometasone [Z= −1.094], and streptozotocin [Z= −1.048]). We found enrichment in the following pharmacological categories: metabolism, ion-transport drugs, ion-transport, and anti-inflammatory drugs.

**Figure 3.**
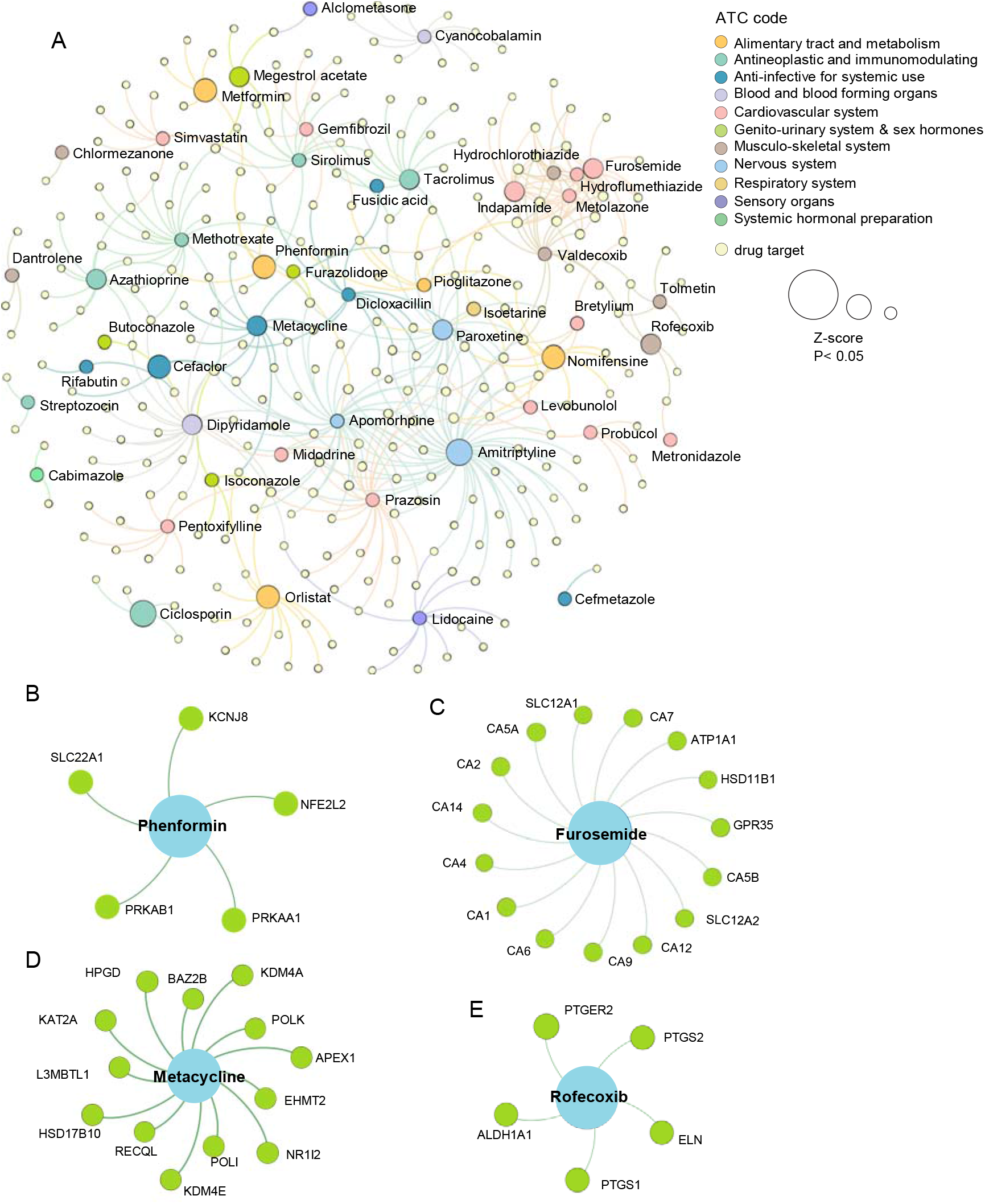
Network-medicine applied to atrial fibrillation drug repurposing. (**A**) A subnetwork is shown to highlight the 54 significant candidate drugs associated with atrial fibrillation (AF) differentially expressed genes and their associated targets. Drugs are colored by their first-level Anatomical Therapeutic Chemical (ATC) classification. (**B-E**) Four candidate drugs with GSEA enrichment score ES>0, P<0.05, and their proposed mechanism of action using our drug-target network analysis.

#### Anti-hypertensive drugs

AF and congestive heart failure (CHF) share risk factors and may represent additive risks for morbidity and mortality. Consequently, both are frequently treated simultaneously. Here, the network-proximity analysis shows a significant association with indapamide (Z= −1.69, P= 0.038, ES= 0.38) and furosemide (Z= −1.91, P= 0.031, ES= 0.38) (**Fig. 3A, C**). Furosemide is an FDA-approved diuretic for the treatment of edema secondary to clinical conditions like CHF by targeting sodium, potassium, chloride transporters (*SLC12A32, SLC12A1*). We found that furosemide targets 13 additional AF module genes (*GPR35, HSD11B1, CA1, CA2, CA4, CA5A, CA5B, CA6, CA7, CA9, CA12, CA14, ATP1A1*) (Figure 3B). Previous studies show that GPR35 contributes to angiotensin II-induced hypertension and cardiotoxicity ^27–29^. Also, AF and CHF are both characterized by abnormal atrial refractoriness, notably from abnormalities in L-type Ca^2+^ current (I_Ca_), transient K^+^ current (I_TO_), slow delayed rectifier current (I_K_), and Na^+^/Ca^2+^ transmembrane exchanger expression and activity ^30^.

#### Anti-inflammatory drugs

Recent evidence suggests inflammation plays a role in AF disease etiology ^31, 32^. Comorbidities associated with AF include low-grade inflammation and metabolic dysfunction disorders, like obesity, hypertension, and coronary artery disease. Furthermore, cardiac muscle and electrical remodeling following AF onset itself can propagate inflammation. Here we identified anti-inflammatory agents rofecoxib (Z= −1.67, P= 0.004, ES= 0.22) and alclometasone (Z= 1.05, P< 0.001, ES= 0.32) as potential AF repurposed drug candidates. Rofecoxib, a COX2 inhibitor FDA approved to treat osteoarthritis and rheumatoid arthritis was shown to target 5 AF module genes (*ALDH1A1, ELN, PTGER2, PTGS1, PTGS2*) (**Figure 3E**).

#### Cardiac metabolism drugs

Dysregulation of cardiac metabolism has been well described in AF^33^. AMPK agonists, including drug family members phenformin and metformin, ranked at the top of AF drug repurposed candidates (Z= −2.475, P=0.002, ES=0.255, and Z=-1.97, P= 0.015, ES=0) (**Fig 3B, Fig 4C**). An association between type II diabetes incidence and AF was established over two decades ago ^34–36^. A prospective case-control analysis of patients treated with metformin demonstrated a decrease of 19% in the incidence of new-onset AF ^37^. AMPK is considered a master regulator of energy status in the heart. Activation of AMPK stimulates pathways that utilize glucose and fatty acids to generate ATP ^38^. In addition to targeting *PRKAB1* (AMPK beta-subunit 1), phenformin also targets a key regulator of oxidative stress, *NFE2L2* (*NRF-2*), as well as the ATP-sensitive potassium channel pore subunit, *KCNJ8*, a metabolically sensitive ion channel and regulator of cardiac repolarization (**Figure 3B**). Further, metformin also targets *ACACB*, *GPD1, ETFDH*, and *DPP4* (**Figure 4C**). Metformin drug target neighbors that were significant and differentially expressed genes in the AF network module or metformin treated a-iCMs included *PTEN, NPPB, GPC2*, and *CXCL12*.

**Figure 4.**
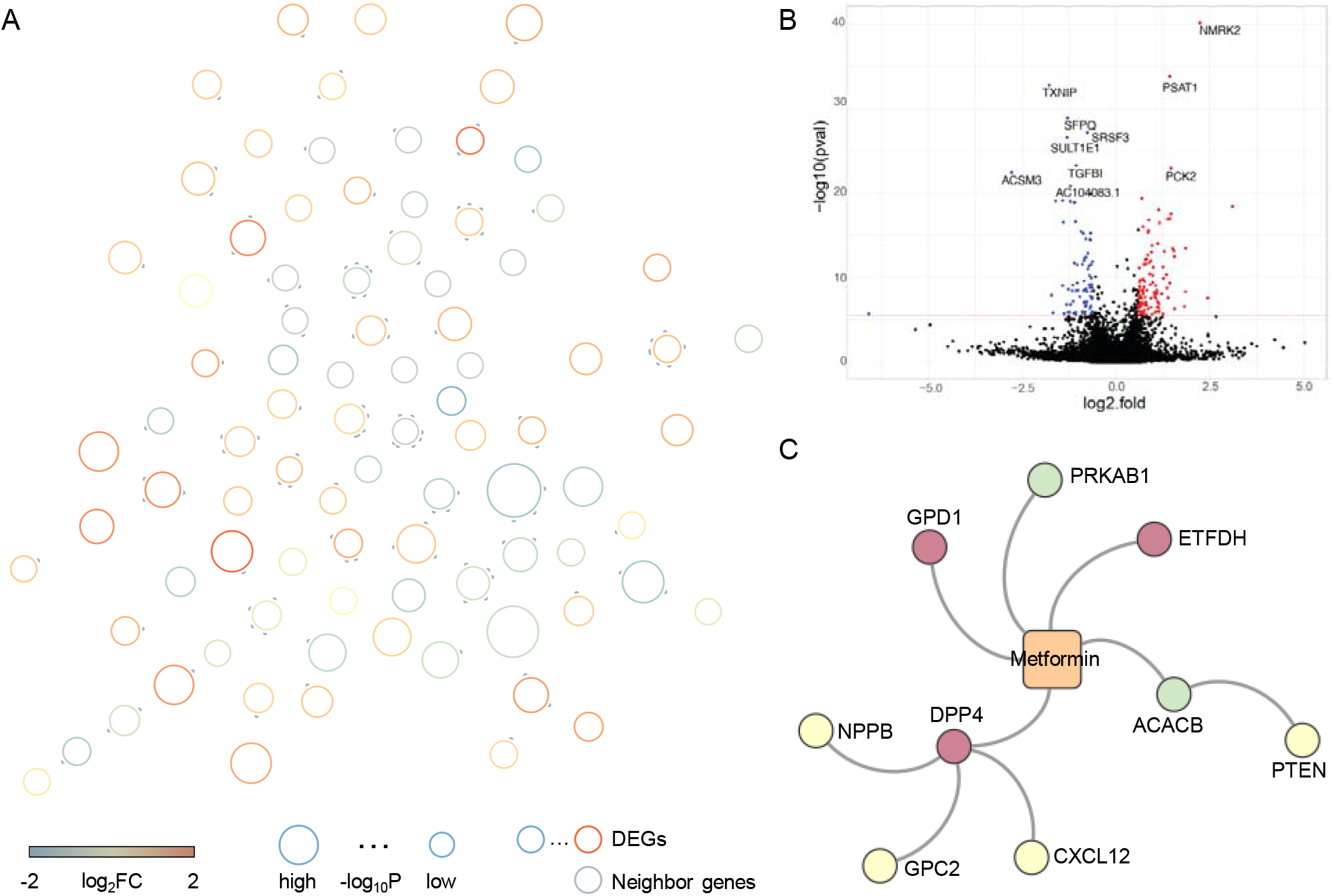
Validation of atrial fibrillation repurposed drug candidate, metformin, using atrial myocytes differentiated from human iPSCs. (**A**) A subnetwork representing the significant differentially expressed (DE) genes and their protein-protein interaction neighbors (gray). The node size is proportional to -log_10_P-value, and color corresponds to log_2_ fold change (log_2_FC) in metformin treated atrial-like cardiomyocytes (a-iCMs) versus control following 1Hz pacing stimulation. (**B**) DE genes of a-iCMs treated with metformin for 48 hours. The red dotted line represents P-value cutoff, P<2.6E-6. Blue and red dots represent genes with decreased and increased expression respectively. (**C**) A subnetwork of netformin, drug targets, and protein-protein interaction neighbors.

### Human iPSC-derived a-iCM validation of Metformin

To validate our *in-silico* drug repurposed findings, we generated atrial myocytes differentiated from human iPSCs (a-iCMs). The iPSC line used in this experiment was obtained from American Type Culture Collection (ATCC, ACS-1030). The iPSCs were reprogrammed from bone marrow derived CD34+ cells obtained from a healthy 31-year-old white female donor using Sendai viral expression of *OCT4, SOX2, KLF4*, and *MYC*. The iPSCs were differentiated to a-iCMs using a-iCMs used for downstream analysis exhibited spontaneous beating and expressed high troponin levels. At 30 days from differentiation onset, human a-iCMs were exposed to either metformin (0.5 mM) or ultra-pure water for 48 hours (**Figure 1C**) with and without 1 Hz pacing stimulation. Beating cells were pretreated ±0.5mM metformin for 6 hrs, then paced for 24 hrs, stimulating a degree of cell stress. All experiment combinations had 3 replicates. Rt-PCR was performed to validate the expression of AF PPI neighbors. With metformin, SCN5A (a PPI partner with AF gene product CAMK2D) was highly expressed in unpaced compared to paced controls. Other known markers associated with low expression in AF increased following metformin treatment in pacing cells (CACNA1C, CACNA1D, HSF1, HSF2, and HSPB1 (HSP27) (**Table 2**). These findings support the favorable effects on proteostasis and ion channel expression.

**Table 2.** Gene expression by rtPCR in unpaced and paced controls a-iCMS following treatment with metformin and control.

|  | Baseline |  | 1 Hz Stimulation |  |
| --- | --- | --- | --- | --- |
|  | Control | Metformin | Control | Metformin |
| HSF1 | 1.02±0.25 | 1.82±0.34 | 1.04±0.31 | 2.95±1.20 |
| HSF2 | 1.00±0.09 | 1.56±0.41 | 1.51±0.70 | 4.72±3.73 |
| HSP27 | 0.02±0.001 | 1.44±0.21 | 1.02±0.21 | 2.03±0.99 |
| SCN5A | 1.03±0.30 | 2.24±0.89 | 1.02±0.24 | 3.44±2.32 |
| CACNA1C | 1.01±0.17 | 1.36±0.58 | 1.00±0.08 | 1.90±0.59 |
| CACNA1D | 1.02±0.24 | 2.21±1.08 | 1.01±0.16 | 2.73±1.15 |

Bulk RNA-seq was used to broadly assess the differentially expressed genes (DEG) associated with metformin exposure (**Fig. 4B**) using a similar design as the rtPCR study (3 replicates per group). Following multiple-testing correction (Bonferroni method with 16,895 expressed genes and family-wise error rate of 0.05), 251 genes exhibited significant differential expression (marginal DEG P < 2.96E-06) for metformin versus water exposure in the context of 1 Hz pacing stimulation. Several key cardiovascular or drug-related markers were significantly upregulated with pacing, such as *NMRK2, PSAT1, PCK2, HMOX1*. KEGG enrichment analysis suggests that the top upregulated DEGs *PSAT1, NMRK2*, and *PCK2* had neighbors enriched in the AMPK/MAPK signaling pathways, metabolism, and inflammation, FoxO signaling, and ion channel regulation (**Figure 5**). *TGFB1* and *TXNIP* were significantly downregulated, indicating enhanced control of redox state and cellular homeostasis. Specifically, we find a subset of known PPI neighbors of TXNIP corresponding to insulin resistance (RELA, SLC2A1, PTPN11) (**Figure 5**). We also find significant downregulation of splicing factors *SFRS3* and *SFPQ. ACSM3*, a rate-limiting enzyme involved in fatty acid metabolism, is significantly downregulated, suggesting a potential decrease in lipid utilization. In summary, these functional observations in a-iCMs mechanistically support that network-predicted metformin is a potential treatment for AF.

**Figure 5.**
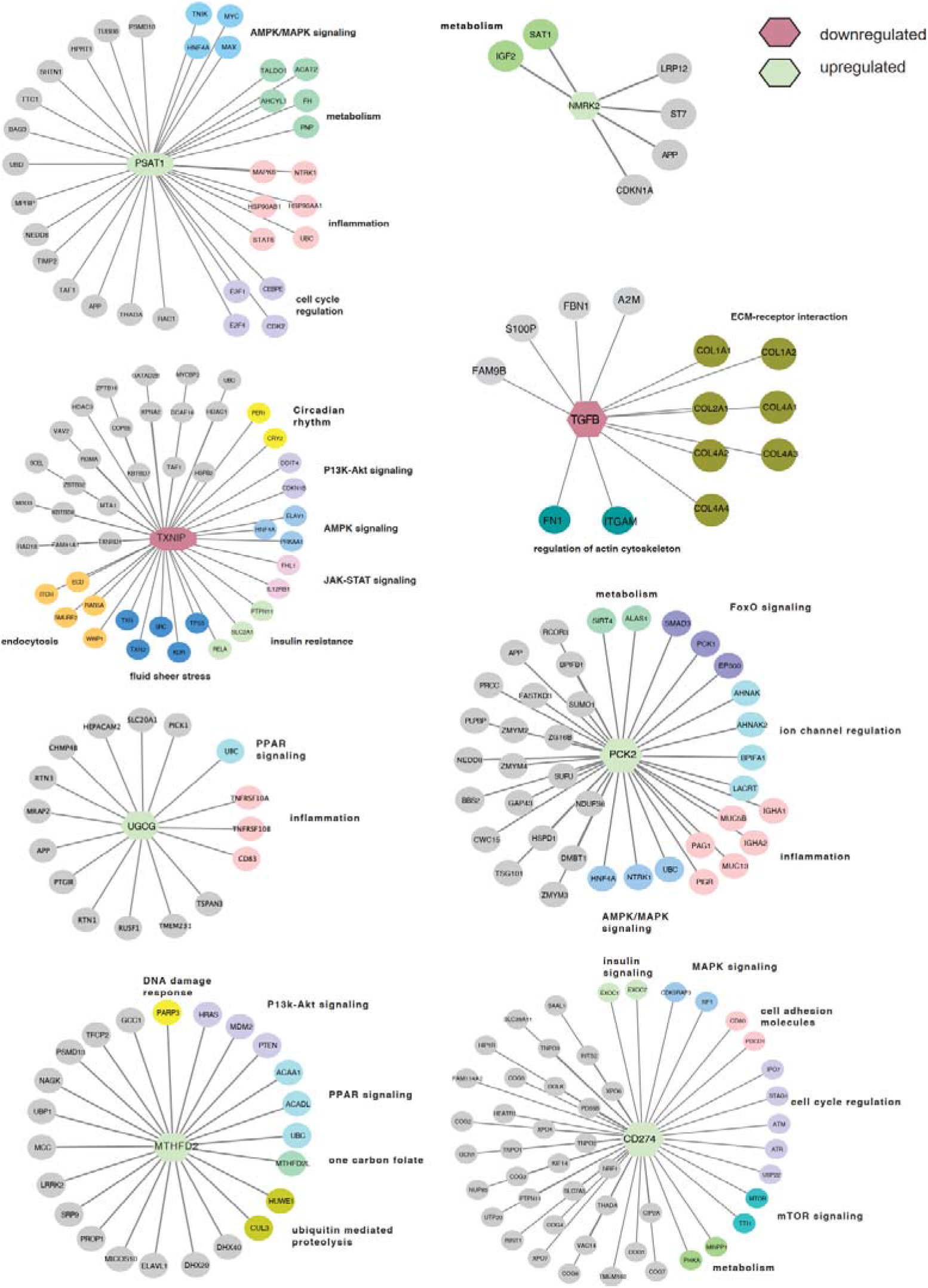
Subnetworks of upregulated and downregulated genes following metformin treatment in a-iCMs. The central node color corresponds to upregulated (green) or downregulated (red) genes following metformin treatment in pacing a-iCMS. Outer node colors correspond to KEGG pathway classification, class is also listed next to each pathway cluster.

## Discussion

Clinical management of AF is limited by high toxicity rates of anti-arrhythmic drugs and poor long-term efficacy. Here, we have generated an AF disease network module and utilized a network-based approach to prioritize alternative drug options for AF. We have shown that our mechanism-based protein-protein interactome can successfully identify drug-disease associations. Specifically, we have identified metformin as a high-quality candidate for drug repurposing for AF using network-proximity analysis of drug targets to the AF gene network.

Metformin is a first-line FDA-approved medication for type II diabetes mellitus. Several studies have provided strong evidence for the association of AF to metabolic syndrome diseases. The Framingham study was one of the first studies showing a significant risk of developing AF in patients with diabetes (OR= 1.4 for men and 1.6 for women, respectively) after multivariable adjustment, and as congestive heart failure and heart valve disease ^34^. Several follow-up studies have strengthened this observation, showing an increased AF risk with longer disease duration or worse glycemic control ^35, 36^. This evidence suggests that in addition to irregularities in conductance control, managing metabolic and inflammatory pathways may provide therapeutic benefits. Metformin targets and activates AMPK, a master regulator of the metabolic stress response, upon sensing high AMP/ATP levels to enhance ATP production and reduce ATP expenditure ^39^. AMPK regulates glucose metabolism and energy utilization, as well as mTORC1 to control autophagy. Additionally, metformin suppresses DRP-1 mediated mitochondrial fission via an AMPK mechanism, reducing mitochondrial fragmentation in mice ^40^, promoting mitophagy to clear dysfunctional mitochondria ^41^, and reducing endoplasmic reticulum (ER) stress, reactive oxygen species production, and protein synthesis, which may reduce proteotoxic stress. Isoproterenol suppresses AMPK that can lead to cardiomyocyte apoptosis and ER stress, and metformin protects against this ^42^.

It is challenging to assess whether metformin’s benefit is owed to it’s drug targets or pleiotropic benefits from weight loss. Indeed, metformin is reported to show a modest decrease in body weight of ~5 kg by reducing food intake ^43^. It improves insulin resistance and inflammation/oxidative stress response mediated by free fatty acids, leptin, and other adipokines, which may target the pathophysiological link between obesity and AF. In the Diabetes Prevention Program, metformin produced durable weight loss for at least 10 yrs of treatment ^44^. In 33 non-diabetic subjects with congestive heart failure, metformin use was associated with improved fasting insulin resistance, reduced weight, and proinflammatory cytokines, including the aging-associated cytokine CCL11 ^45^. A meta-analysis showed that metformin use in persons at risk for DM improved weight, lipids, and insulin resistance and reduced new-onset DM by 40% ^46^.

In an observational study of 645,710 T2DM subjects over 13-year follow-up, metformin use was associated with 20% less AF ^47^. However, there are no prospective trials reported of metformin for AF in non-diabetics. Metformin has been proposed as upstream therapy in patients scheduled for AF ablation (clinicaltrials.gov NCT02931253). We aim to perform the first study to assess the effect of metformin for the prevention of AF progression in non-diabetics. Prior evidence has alluded to the therapeutic benefits of metformin for AF. An ongoing clinical trial, Upstream Targeting for the Prevention of Atrial Fibrillation: Targeting Risk Interventions and Metformin for Atrial Fibrillation (TRIM-AF) is investigating the benefit of metformin and lifestyle/risk factor modification interventions in AF patients (NCT 03603912). Both interventions have been reported to target AMP-activated protein kinase (AMPK), a heterodimeric protein composed of alpha (catalytic) and beta/gamma (regulatory) subunits, encoded by genes including PRKAA1, PRKAA2, and PRKAB1. Of note, PRKAA1 was present in the AF subnetwork.

In addition to metformin, several inflammation regulating therapies were identified using our approach, including rofecoxib and alclometasone. Localized and systemic inflammation have both been described to coincide with AF. Circulating inflammatory biomarkers that have been associated with incident AF include C-reactive protein (CRP), tumor necrosis factor (TNF), IL-6, and NF-KB ^48^. Analysis of 5,806 subjects in the Cardiovascular Health Study showed that CRP levels above the median predicted an increased risk of developing AF ^49^. Another study showed that circulating CRP levels are increased in patients with AF compared to those in normal sinus rhythm ^50^.

We found TGFB1 as a significant DEG node in our AF network module, which was downregulated following metformin treatment in the iCMs. In clinical studies of patients with AF compared to patients with normal sinus rhythm, serum concentrations of TNF-alpha and TGFB1 were increased ^51, 52^. TGFB1 is a well-known pro-fibrotic cytokine that promotes atrial structural and electrical remodeling of the atria ^53^. Interstitial fibrosis promotes slow heterogeneous electrical conduction between myocytes, contributing to a substrate for reentrant electrical activity. TGFB1 may also affect calcium handling. Reduced L-type calcium channel current (I_CaL_) and reduced expression of Ca_v_1.2 (CACNA1C) following TGFB1 exposure have been reported in neonatal rat atrial myocytes, but not yet been reported in human atrial tissues ^54^. Suppl. Fig 3 shows an inverse relation of TGFB1 with CACNA1C in adult human left atrial appendage tissues (n=265, P<0.001).

Interestingly, we found NPPB (brain natriuretic peptide, BNP) and CXCL12 as PPI neighbors of the metformin target, DPP4 (**Figure 4C**). BNP can suppress the activity of the renin-aldosterone-angiotensin system (RAAS) ^55^. Overstimulating the RAAS via angiotensin II has been clinically shown to promotes localized oxidative stress by activating NF-κB and increasing the production of IL-6 and CRP. As noted above, these inflammatory markers are elevated in patients with AF rhythm. The cytokine CXCL12, also a PPI neighbor of DPP4, is responsible for recruiting monocytes and lymphocytes. Systemic levels of macrophages and activated T lymphocytes are elevated in patients with persistent AF ^56, 57^. These findings strengthen the powerful role inflammation plays in AF disease etiology.

We acknowledged several potential limitations in the current study. For example, incompleteness of the human protein-protein interactome and drug-target networks may influence the model performance. Disease heterogeneities of bulk transcriptomics data may influence the AF disease network module analysis as well. Integration of single-cell transcriptomics may help to better understand disease heterogeneities of AF and identify cell type-specific drug targets and treatment approaches for AF in the near future. In addition, integration of AF genetics findings may help to identify more likely risk genes and drug targets to accelerate effective therapeutic discovery in AF as well.

In summary, we have identified AF specific dysregulated networks enriched in cardiac metabolism, ion transport, and immune pathways. Importantly, we utilized network-medicine strategies to identify 9 candidate drugs that are putative repurposed drugs for AF. Utilizing multiple lines of evidence, we nominated metformin for functional validation and identified key molecular signals that help explain metformin’s mechanism of action. We believe that the network medicine approaches presented here, if broadly applied, would significantly catalyze effective treatment development in AF and other cardiovascular diseases as well.

## Materials & Methods

### Human Induced Pluripotent Stem Cell Derived atrial cardiomyocytes

These studies used a line of inducible pluripotent stem cells (iPSC), obtained from a female patient (ATCC, ACS-1030). iPSCs are routinely used between passage 20-70. Pluripotency to form atrial-like cardiomyocytes (a-iCMs) was confirmed routinely by assessment of pluripotency genes (TRA-160, SSEA-4) and ability to differentiate and create beating cardiomyocytes. iPSCs were grown in E8 media, passaged every 3 days, and incubated at 37°C, 5% CO_2_, 98% relative humidity ^58^. Differentiation of a-iCMs was adapted from the protocol of Burridge and colleagues, with the inclusion of retinoic acid (1 μmol/L) in the differentiation media from days 4-8 to enhance atrial-like gene expression ^58–60^. A metabolic selection step (no glucose, 4 mM lactate) was used from days 12-14 to limit the abundance of non-myocytes in the cultures. The media was returned to CDM3 from day 16 until day 20, followed by maintenance in CDM3-M media until the cells were used for experiments. Using this method, beating cells were typically evident by day 10, and beating sheets (electrically synchronized cell sheets covering the entire dish) by day 20. One week prior to the stimulation experiment, cells were dissociated, combined and replated at a density of 150,000 per well in 12 well dishes. Experiments began after differentiation for 30 days and were completed in triplicate.

### Drug Treatment

a-iCMs were pre-incubated for 6 hours using 0.5 mM metformin in water prior to electrical pacing for 24 hours. Cells in the presence of water only were used as controls.

### RNA isolation and sequencing

RNA was isolated from the a-iCMs (1.2 million cells) following the RNeasy Plus protocol ^61–63^. RNA was eluted in 15 ul of RNase-free water, with concentration determined by OD_260_ nm and RNA stored at −80°C. Libraries were constructed using lllumina oligo-dT based stranded kits following the recommended protocol. Approximately 60 million 100-basepair paired ends reads (30 million clusters) were generated for each library on an Illumina NovaSeq 6000 sequencer with a S1 flowcell. Both library construction and sequencing were done at the University of Chicago Genomics Facility. Tissue processing, RNA isolation and sequencing from the human atrial tissues was described in a prior publication ^21^.

### Building the human protein-protein interactome

To build a comprehensive list of human protein-protein interactions, we assembled data from 18 bioinformatics and systems biology databases compromised of five experimental assays: (i) binary PPIs tested by high-throughput yeast-two-hybrid (Y2H) systems; (ii) binary, physical PPIs from protein 3D structures; (iii) kinase-substrate interactions by literature-derived low throughput or high-throughput assays; (iv) signaling network by literature-derived low-throughput experiments; and (v) literature-curated PPIs identified by affinity purification followed by mass spectrometry (AP-MS), Y2H, or by literature-derived low-throughput assays. The genes were mapped to their ENTREZ ID based on the NCBI database^64^ as well as their official gene symbols on GeneCards (https://www.genecards.org/). The human protein-protein interactome used in this study includes 351,444 unique PPIs (edges or links) connecting 17,706 proteins (nodes).

### Differential Gene Expression Analysis from a-iCM RNAseq

RNAseq results from 6 wells of a-iCMs in the metformin versus water study were quality accessed using the nf-core/rnaseq pipeline ^65^. Transcript-level quantifications per sample were generated using kallisto with the Gencode version 31 human transcriptome ^66^. Gene-level differential expression p-values were obtained from transcript-level differential expression p-values using sleuth and the Lancaster combining method ^67, 68^.

### Human subjects

Left atrial appendage (LAA) tissues were collected from 251 patients who underwent elective cardiac surgery to treat AF, valve disease, or other cardiac disorders, and from 14 non-failing donor atria. The study was approved by the Cleveland Clinic Institution Review Board (IRB). Further details have been described in prior reports ^21^.

### Differential Gene Expression Analysis from Human Left Atrial Tissue RNASeq

After preprocessing and quality control, transcript-level quantifications per sample were generated using kallisto with the Gencode version 27 human transcriptome ^66^. Genelevel quantifications were generated from these transcript-level estimates using the tximport package ^69^ with the length scaled TPM option; after sample and gene expression level filtering, 19,816 genes measured on 280 RNAseq profiles from (distinct patient) left atrial appendage tissues were used for DGE analyses. A description of the corresponding patient cohort was previously reported ^21^. Linear regression models per gene with empirical Bayes shrinkage ^70^ as implemented in the limma R package were used to estimate and test the contrast between those whose hearts were in atrial fibrillation versus sinus rhythm at time of heart tissue acquisition (surgery or donor heart harvesting). The per-gene regression models included adjustments for sex, age (modeled as a 2 degree of freedom spline), white race, interactions of sex with age and sex with race, as well as 33 surrogate variables (SV) to help account for unmodeled confounders and heterogeneity. SVs were estimated using a latent high dimensional confounder approach ^71^ as implemented in the cate R package.

### Building the Atrial Fibrillation disease network

We used the DEGs from the human left atrial appendage RNA-seq data. We extracted the PPIs for the DEGs from the human interactome.

### Pathway enrichment analysis

Gene Ontology (GO) and KEGG pathway enrichment analysis was used to identify pathways enriched in the AF disease module and Metformin treated hiPSC-CM differentially expressed genes using the enrichR R package (version 3.0) ^72^.

### Building the drug-target network

We collected drug-target interaction information from DrugBank database (accessed April 2021) ^73^, Therapeutic Target Database (TTD) ^74^. Drug-target interactions meeting the following three criteria used: (i) binding affinities, including K_i_, K_d_, IC_50_, or EC_50_ each ≤ 10 μM; (ii) the target was marked as “reviewed” in the UniProt database (ref); and (iii) the human target was represented by a unique UniProt accession number.

### Network proximity measure

To quantify the relationship of the AF disease network and Metformin treated hIPSC-CMs DEGs, we adopted the closest path-based network proximity measure as below.

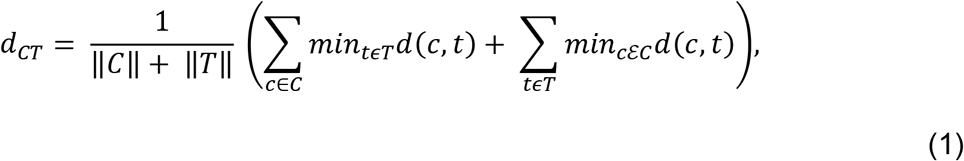

where *d*(*c,t*) is the shortest distance between gene *c* and *t* in the human protein interactome. The network proximity was normalized to Z-score based on permutation tests:

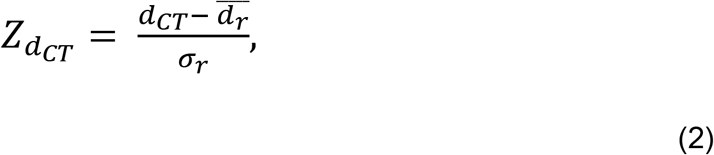

Where 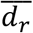 and *σ_r_* were the mean and standard deviation of the permutation test was repeated 1000 times using two randomly selected gene lists with similar degree distributions to those of *C* and *T*. The *P* value was calculated based on the permutation test results. *Z*-score < −1.0 and *P* < 0.05 were considered significant proximal AF disease associations. The networks were visualized using Gephi 0.9.2 (https://gephi.org/) or Cytoscape 3.8.2 (https://cytoscape.org/).

### Gene set enrichment analysis

To further prioritize drugs identified by the drug-gene network analysis, we performed gene set enrichment analysis. Differential expressed genes from the AF disease module with P < 0.05 were used, as well as differential gene expression in cells treated with various drugs downloaded from the Connectivity Map (CMAP) database ^26^. For each drug that was in both the CMAP data set and our drug-target network, we calculated an enrichment score (ES) based on previously described methods ^75^ as shown below:

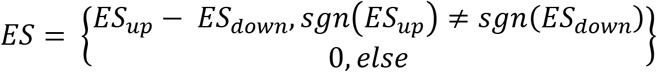

ES_up_ and ES_down_ were calculated for up- and down-regulated genes from the AF disease network signature. We first computed a_up/down_ and b_up/down_ using the following equations:

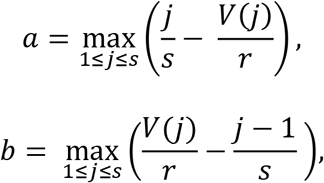

Where *j* = 1, 2, 3…., *s* corresponds to the genes of the AF disease signature in ascending ranked order. *V*(*j*) corresponds to the rank of gene *j*, where 1 ≤ *V*(*j*) ≤ *r*, with r being the number of genes (12,849) from the drug profile. *ES _up/down_* were defined by:

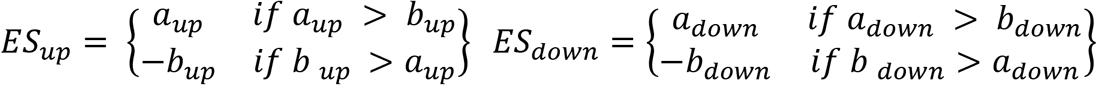

Permutation tests were repeated 100 times using a random gene list with the same up- and down-regulated genes as the AF signature dataset used to measure the significance of the ES scores. Drugs were considered to have potential treatment effect if ES > 0 and P<0.05.

## Data and Code Availability

The codes written for and data used for network proximity analysis are available from website: https://github.com/ChengF-Lab/

## Authors Contributors

F.C., M.K.C. and D.R.V. conceived the study. J.C.L., Y.Z., S.R.G, and J.H.R. performed all experiments and data analysis. J.B. contributed to differential expression analysis and helped to interpret the data analysis. J.C.L., F.C., D.R.V. and M.K.C. drafted the manuscript and critically revised the manuscript. All authors critically revised and gave final approval of the manuscript.

---

**Suppl. Figure 1.**
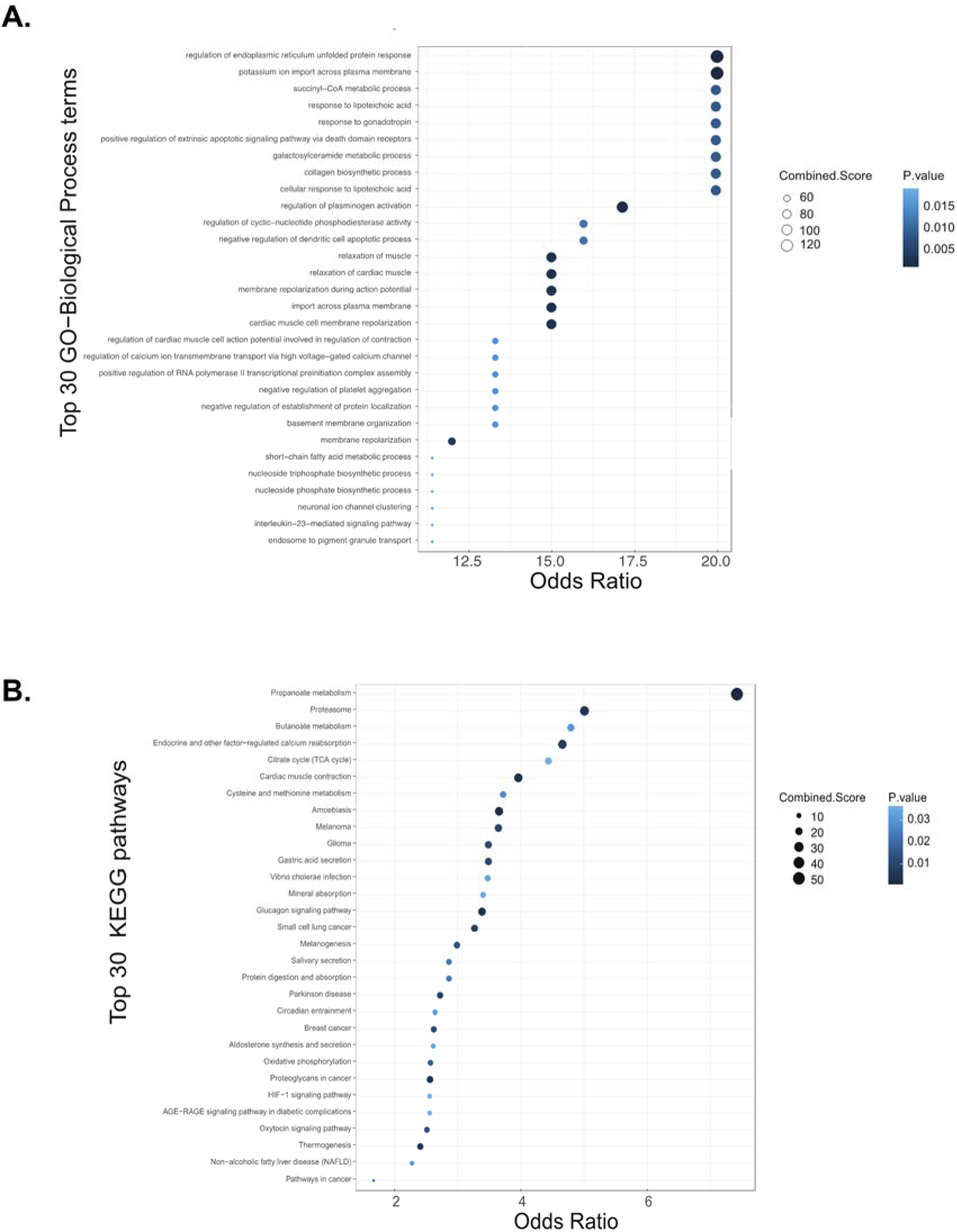
Pathway enrichment analysis of the top genes enriched in the AF disease network. **A.** Gene ontology (GO) expression analysis. **C.** KEGG pathway enrichment analysis. The node size corresponds to combined score, and color corresponds to P-value.

**Suppl. Figure 2.**
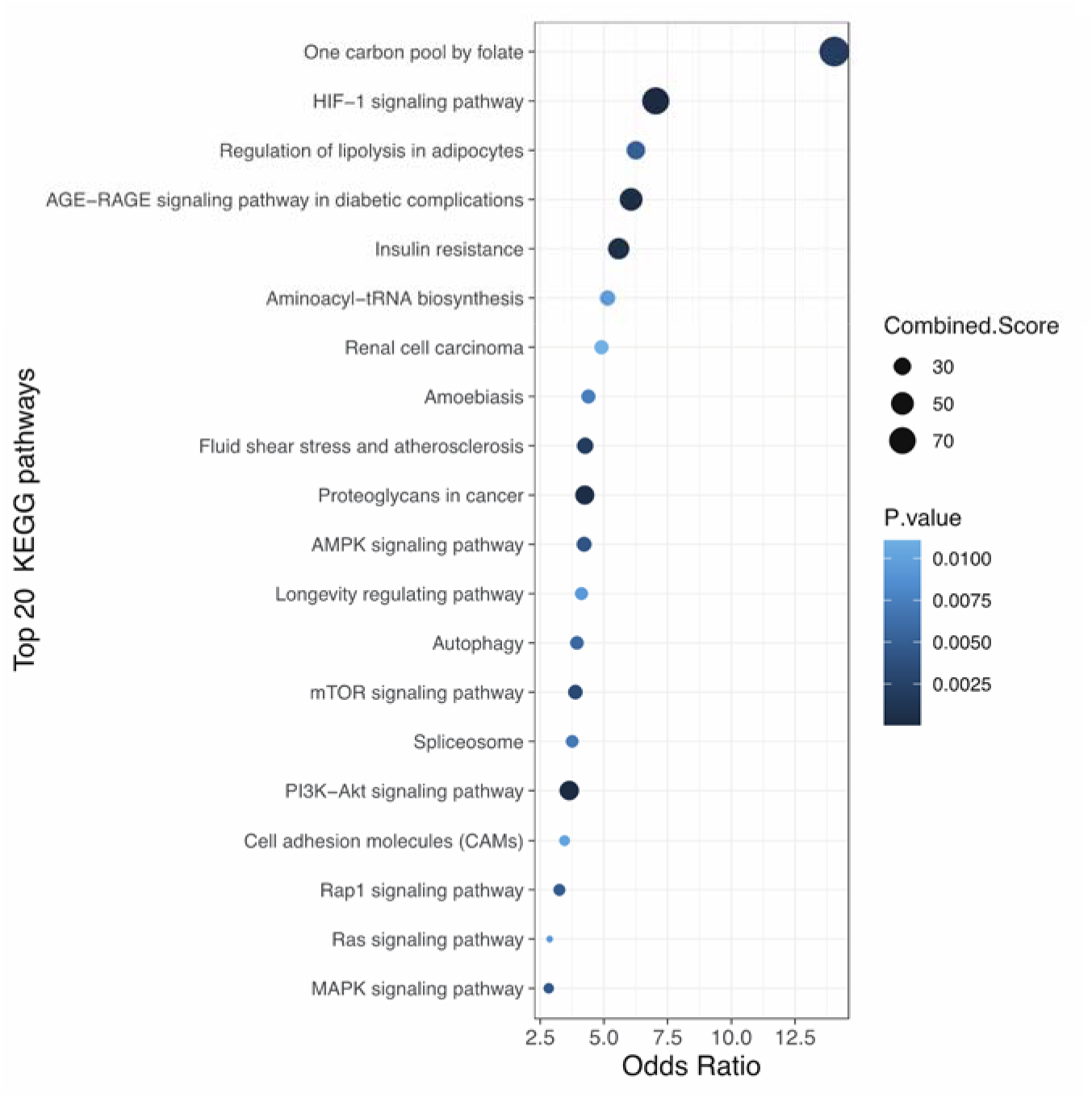
Pathway enrichment analysis of the top genes enriched in pacing a-iCMS following metformin treatment. **A**. Gene ontology (GO) expression analysis. **C.** KEGG pathway enrichment analysis. The node size corresponds to combined score, and color corresponds to P-value.

**Suppl. Figure 3.**
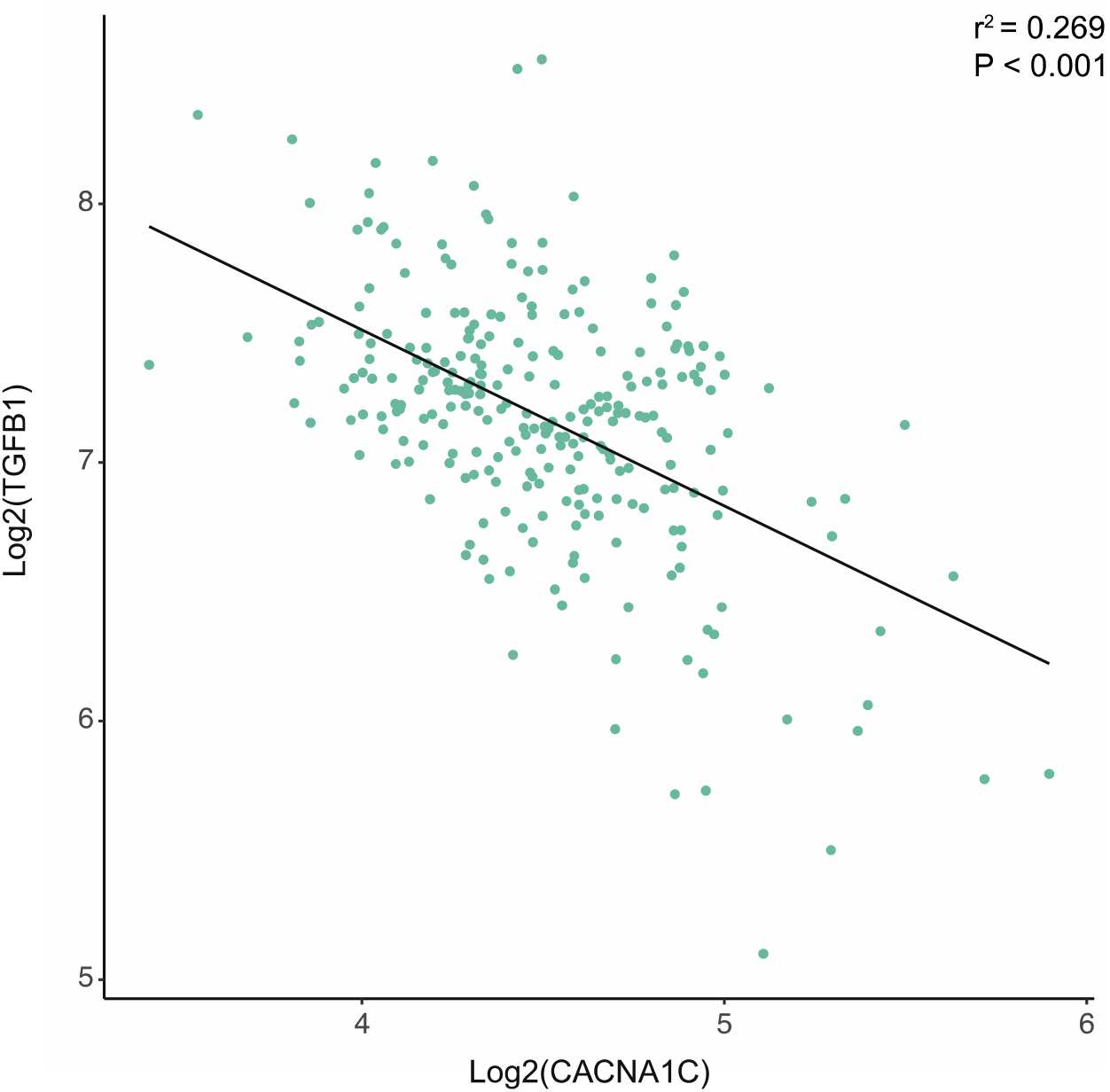
RNAseq analysis documents an inverse relationship between TGFB1 and CACNA1C abundance in human left atrial appendages.

